# Painting cell-cell interactions by horseradish peroxidase and endogenously generated hydrogen peroxide

**DOI:** 10.1101/2024.06.11.598589

**Authors:** Inyoung Jeong, Kwang-eun Kim, Hyun-Woo Rhee

**Author notes:** **Corresponding Author Kwang-eun Kim** − Department of Chemistry, Seoul National University, Seoul 08826, Korea; Department of Convergence Medicine, Yonsei University Wonju College of Medicine, Wonju 26426, Korea;, **Hyun-Woo Rhee** − Department of Chemistry, Seoul National University, Seoul 08826, Korea.

## Abstract

Cell-Cell interactions are fundamental in biology for maintaining physiological conditions, with direct contact being the most straightforward mode of interaction. Recent advancements have led to the development of various chemical tools for detecting or identifying these interactions. However, the use of exogenous cues, such as toxic reagents, bulky probes, and light irradiations, can disrupt normal cell physiology. For example, the toxicity of hydrogen peroxide (H_2_O_2_) limits the applications of peroxidases in proximity labeling field. In this study, we aimed to address this limitation by demonstrating that membrane-localized Horseradish Peroxidase (HRP-TM) efficiently utilizes endogenously generated extracellular H_2_O_2_. By harnessing endogenous H_2_O_2_, we observed that HRP-TM-expressing cells can effectively label contacting cells without the need for exogenous H_2_O_2_ treatment. Furthermore, we confirmed that HRP-TM labels proximal cells in an interaction-dependent manner. These findings offer a novel approach for studying cell-cell interactions under more physiological conditions, without the confounding effects of exogenous stimuli. Our study contributes to elucidating cell-cell interaction networks in various model organisms, providing valuable insights into the dynamic interplay between cells in their native network.

---

Cell-Cell interactions are crucial for regulating cellular physiology, playing essential roles in neuronal signaling, tissue formation, immune responses, and tumor progression ^[1-4]^. T cells, for instance, are activated when they recognize antigen-presenting cells by direct contact, while tumor microenvironments consist of interactions among various cell types, including cancer cells and non-cancerous cells such as immune cells, endothelial cells, or fibroblasts ^[5]^. It is essential to identify intercellular networks at the tissue level while maintaining physiological conditions.

Recently, several studies on cell-cell interaction have been reported with diverse approaches. One of the most widely used is a fluorescence-based method. Porterfield et al. developed a tool to detect cell-cell contact via luciferase-luciferin reaction ^[6]^. In addition, Tang et al. suggested a new tool, G-baToN, in which GFP and anti-GFP nanobody (αGFP) record the intercellular interactions ^[7]^. Another method based on a cell-penetrating fluorescent protein was suggested by Ombrato et al., as a system for identifying the spatial cellular environment of metastatic cancers ^[8]^. However, the above methods require engineered ligand and receptor expression, which makes it impossible to reveal cell-cell interactions between unknown cell types, and the interaction between artificial ligand-receptor pair could influence physiological cell-cell contact events.

Proximity labeling has become a versatile tool for studying spatiotemporal proteomics and has emerged as a powerful tool for deciphering cell-cell interaction ^[1]^. LIPSTIC, which utilizes sortaseA (SrtA) to transfer LPXTG-biotin to N-terminal oligoglycine of proteins at the cell surface, confirmed the interactions of dendritic cells and T-cells ^[9]^. Recently, it has been developed into EXCELL ^[10]^, an engineered SrtA (mgSrtA) to functionalize the enzyme for transferring LPXTG-biotin to N-terminal monoglycine. However, the relatively bulky conjugate (LPETG=520 Da) compared to biotin (244 Da) makes the tissue penetration of LPETG-biotin uncertain. PUP-IT ^[11]^ and FucoID ^[12, 13]^ have also been reported as methods that allow exogenous enzymes to be expressed in bait cells, enabling protein/peptide or glycosylation probes to be displayed on prey cells. In this case, direct cell-cell interactions can be selectively detected by enzymatic reaction, however, efficient delivery of large molecular weight probes (e.g. LPETG peptide, PUP protein, fucosylation donor) for these proximity labeling methods should be optimized for *in vivo* application.

Recently, photocatalysts have been developed to capture spatial biological network or spatiome ^[14]^. MicroMap (μMap), a photocatalytic reaction of Iridium that generates reactive carbene species upon blue light irradiation, has been suggested as a tool with a significantly short labeling radius of around 4 nm ^[15]^. Oslund et al. have designed riboflavin tetraacetate (RFT) mediated labeling of proteins, called photocatalytic cell-tagging (PhoTag) ^[16]^. Lie et al. reported antigen-specific T cell detection using the photosensitizer dibromofluorescein (DBF) termed PhoXCELL ^[17]^, and Qiu et al. developed Ru-^1^O_2_-hydrazide system for photocatalytic cell labeling ^[18]^. However, light irradiation for photocatalytic reactions critically limits their use to *in vitro* or cell-level applications rather than *in vivo*.

Owing to the aforementioned potential limitations of proximity labeling tools for *in vivo* application for labeling cell-cell interaction, we hypothesized that peroxidases could effectively address these challenges. Peroxidases such as APEX2 and Horseradish Peroxidase (HRP) catalyze the oxidation of phenol substrates like desthiobiotin-phenol (DBP= 333.43 Da) ^[19]^ to generate phenoxy radicals in the presence of hydrogen peroxide (H_2_O_2_). This reaction enables protein labeling by forming covalent bonds between biotin-phenoxy radicals and electron-rich amino acid residues, such as tyrosine. Previous studies have utilized exogenous HRP, split HRP, or nucleic acid-based HRP mimics, such as G-quadruplex/hemin, to label and identify surface proteomes and intercellular protein-protein interactions in various *in vitro* or *ex vivo* models ^[20, 21, 22, 23]^. However, the toxicity of exogenous H_2_O_2_ treatment limits its application.

Recent research has demonstrated that live cells generate endogenous H_2_O_2_ under physiological conditions ^[24, 25, 26]^. In the extracellular space, endogenous reactive oxygen species (ROS) is generated by NADPH oxidase (NOX) ^[27]^ or H_2_O_2_ can also diffuse directly across the membrane via aquaporin ^[28]^. Consequently, extracellular H_2_O_2_ is continuously produced through this process. Study by the Hamachi group has reported that endogenously generated H_2_O_2_ can activate chemical probe for proximal protein labeling ^[24]^ and our group also reported that subcellularly expressed APEX2 can be activated by endogenously generated ROS in live cells ^[25]^. Ju and colleagues successfully labeled protein on the cell surface using HRP, utilizing endogenously generated H_2_O_2_ during cell starvation ^[26]^. Thus, we hypothesized that the APEX2/HRP reaction could utilize H_2_O_2_ on the cell contact under the physiological condition.

To develop a proximity labeling tool for mapping intercellular interactions, we designed an APEX2/HRP construct with Igκ leader signal peptide (SP) in the N-terminus and the transmembrane domain (TM) of PDGF receptor beta, which exposes the peroxidase extracellularly. We anticipated that the peroxidase reaction with DBP, with or without H_2_O_2_ treatment, would biotinylate surface proteins of HRP-TM expressing cells and their proximal cells (**Figure 1**).

**Figure 1.**
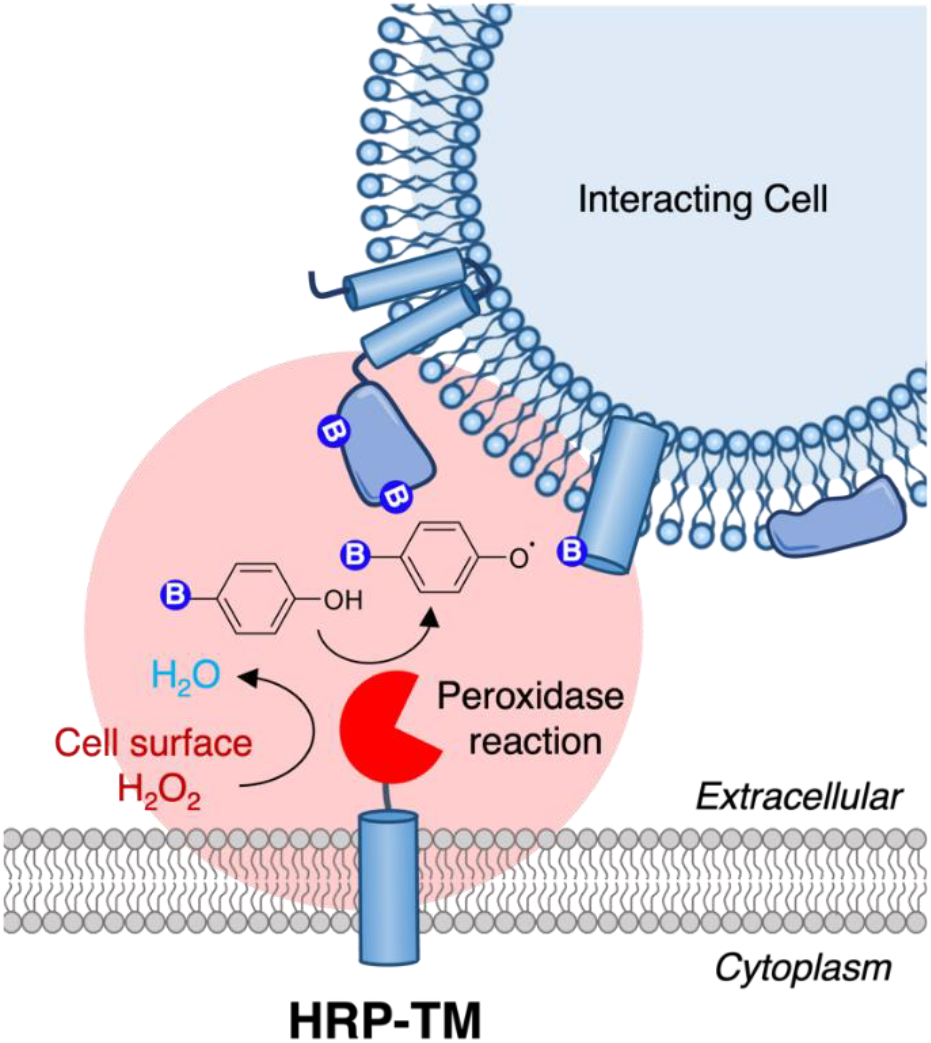
Scheme of endogenous hydrogen peroxide-assisted proximal cell labeling. HRP is expressed in the extracellular space by fusing transmembrane domain (TM). Upon biotin-phenol addition, HRP biotinylates membrane proteins of proximal cells by utilizing endogenous H_2_O_2_ generated in the extracellular space.

## Results

First, we compared the utilization of endogenous H_2_O_2_ between APEX2 and HRP. HRP-TM or APEX2-TM were transiently expressed in HEK293T cells to compare the labeling efficiency with or without exogenous H_2_O_2_ treatment. HRP-TM is fused with a Myc tag, and APEX2-TM is fused with a V5 tag, respectively. Anti-Myc and Anti-V5 blots confirmed the expression of each construct (**Figure 2A**). Non-specific bands in the V5 blot indicated equal protein loading. Biotinylated proteins were detected in the Streptavidin(SA)-HRP blot. With additional H_2_O_2_ treatment, both HRP and APEX2 effectively biotinylated proteins (**Figure 2A**). Immunofluorescence imaging confirmed the labeling of the cell surface under H_2_O_2_ treatment condition (**Figure S1**).

**Figure 2.**
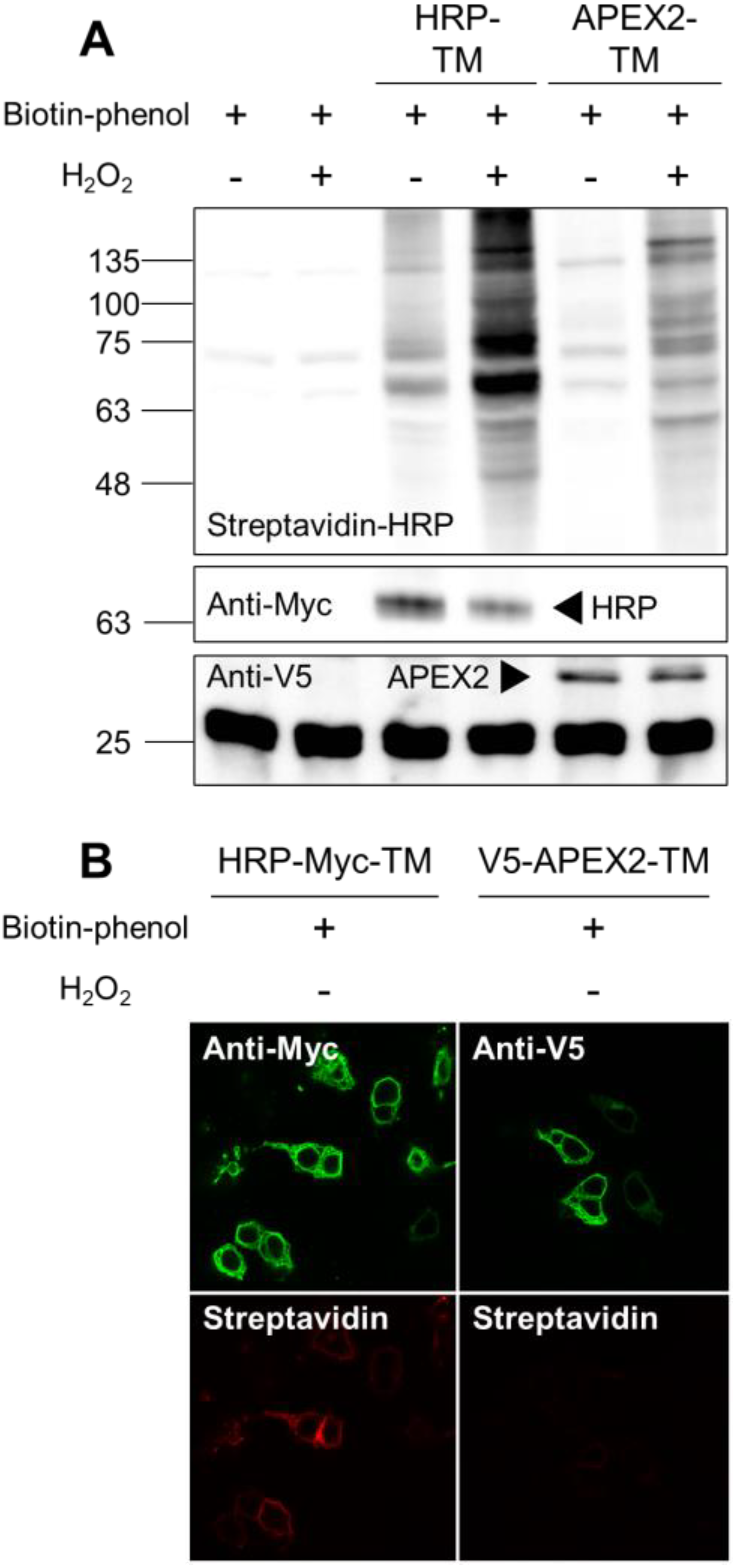
Comparison of H_2_O_2_ facilitating activity of peroxidase at the cell surface. (A) Western blots of biotinylated proteins (SA-HRP) labeled by HRP-TM or APEX2-TM with or without exogenous H_2_O_2_ treatment. (B) Immunofluorescence imaging HRP-TM (Anti-Myc) or APEX2-TM (Anti-V5) and biotinylated proteins (Streptavidin-Alexa).

Interestingly, without H_2_O_2_ treatment, HRP still showed cell surface biotinylation, unlike APEX2 (**Figure 2A, B**). In the imaging experiments, Anti-Myc and Anti-V5 confirmed the cell surface localization of HRP-TM and APEX-TM. Without exogenous H_2_O_2_, cell surface biotinylation was only detected in HRP-TM expressing cells. We observed that HRP exhibited higher labeling efficiency than APEX2. This can be explained by HRP having a 4-fold higher *k*_cat_/K_M_ value than APEX2 ^[29]^, and HRP, but not APEX2, has glycosylation ^[30]^ contributing to effective labeling on the plasma membrane. Our findings suggest the possibility of HRP-TM could label proximal cells utilizing endogenously generated H_2_O_2_.

In this experiment, we also prepared TurboID-TM expressed cells for a comparative experiment. It is noteworthy that TurboID, an engineered biotin ligase has been applied to study astrocyte-neuron interactions at tripartite peri-synapses in the mouse brain. However, TurboID catalyze the labeling reaction from biotin and ATP and it has been pointed out that the extracellular ATP concentration is 1000-fold lower than the estimated K_M_ value of TurboID ^[31]^ and it is expected that labeling intensity of TurboID at the cell surface can be highly dependent of extracellular concentration of ATP which requires experimental validation. As expected, it was shown that cell surface TurboID had significantly lower labeling efficiency than HRP and APEX2, despite the addition of ATP (**Figure S2**) possibly due to the active hydrolase activity for ATP at the cell surface ^[32]^. Since excess treatment of ATP can cause innate immune response by purinergic receptor ^[33]^, we excluded the utilization TurboID in our cell-cell labeling study and we focused utilization of HRP for cell-cell interaction labeling.

To confirm that HRP-TM can label the cell-cell contact site, we generated 293 Flip-In T-REx cell lines that stably express HRP-TM on the cell surface. Wild-type (WT) cells were stained with the green CMFDA dye in advance and co-cultured with HRP-TM expressing cells for 24 hours. Then, the cells were incubated and stained for confocal fluorescence microscopy imaging (**Figure 3A**). When excessive H_2_O_2_ was treated to cells, diffusive labeling was observed (**Figure 3B**) On the other hand, cells labeled only with endogenous H_2_O_2_ showed a defined radius of biotinylation, with only about half of the cell membrane being labeled. This result indicates that HRP-TM could successfully paint proximal cells with DBP in contact by utilizing endogenous H_2_O_2_ and shows more spatially restricted biotinylation on cells (**Figure 3C**).

**Figure 3.**
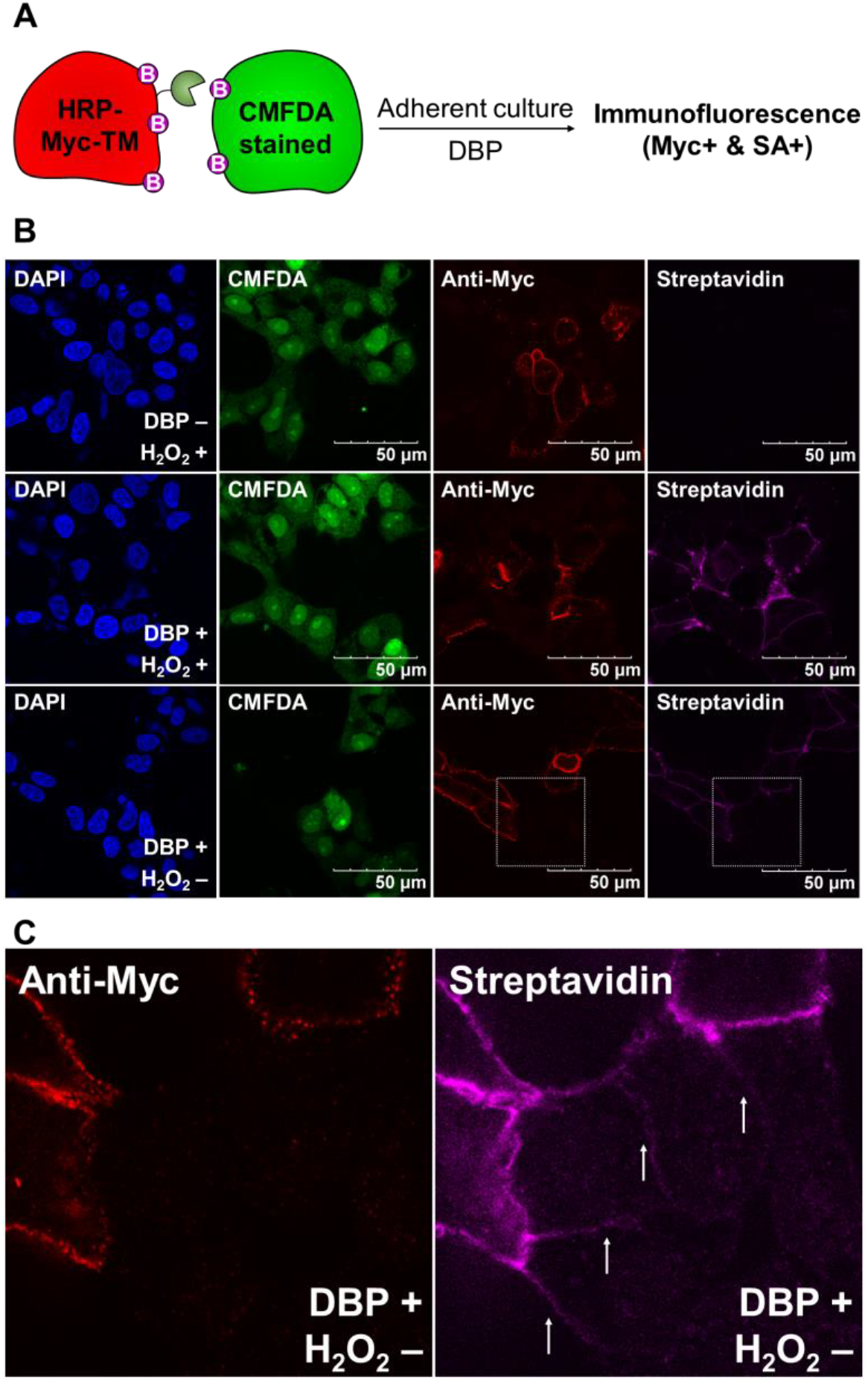
Contact-dependent biotinylation via HRP-TM. (A) HRP-TM stable cells were co-cultured adherently with WT cells stained with green CMFDA dye, and DBP was added for HRP-mediated biotinylation. Cells were stained with DAPI, mouse anti-Myc, anti-mouse-AF568, and Streptavidin-AF647 for fluorescence microscopy. (B) Confocal fluorescence microscopy images of HRP-TM mediated biotinylation of contacting cells. (C) Zoom-in regions in (B).

Next, to test whether HRP-TM can successfully label interacting cells specifically, we employed two pairs of epitope tag-nanobody to induce intercellular interactions: HAFrankenbody (HAFB, 26.6 kDa)-3xHA tag (YPYDVPDYA) ^[34]^ and ALFAnanobody (NbALFA, 14.5 kDa) -ALFA tag (SRLEEELRRRLTE) ^[35]^. HAFB and NbALFA are known to bind their target tags strongly with high binding affinity of K_D_ = 14.7 ± 7.4 nM and K_D_ = ∼26pM respectively ^[34, 35]^. HAFB (Cell A) and HA-tag (Cell a), or NbALFA (Cell B) and ALFA-tag (Cell b) will result in two cells in proximity, thereby biotinylation across cell-cell interface (**Figure 4A**).

**Figure 4.**
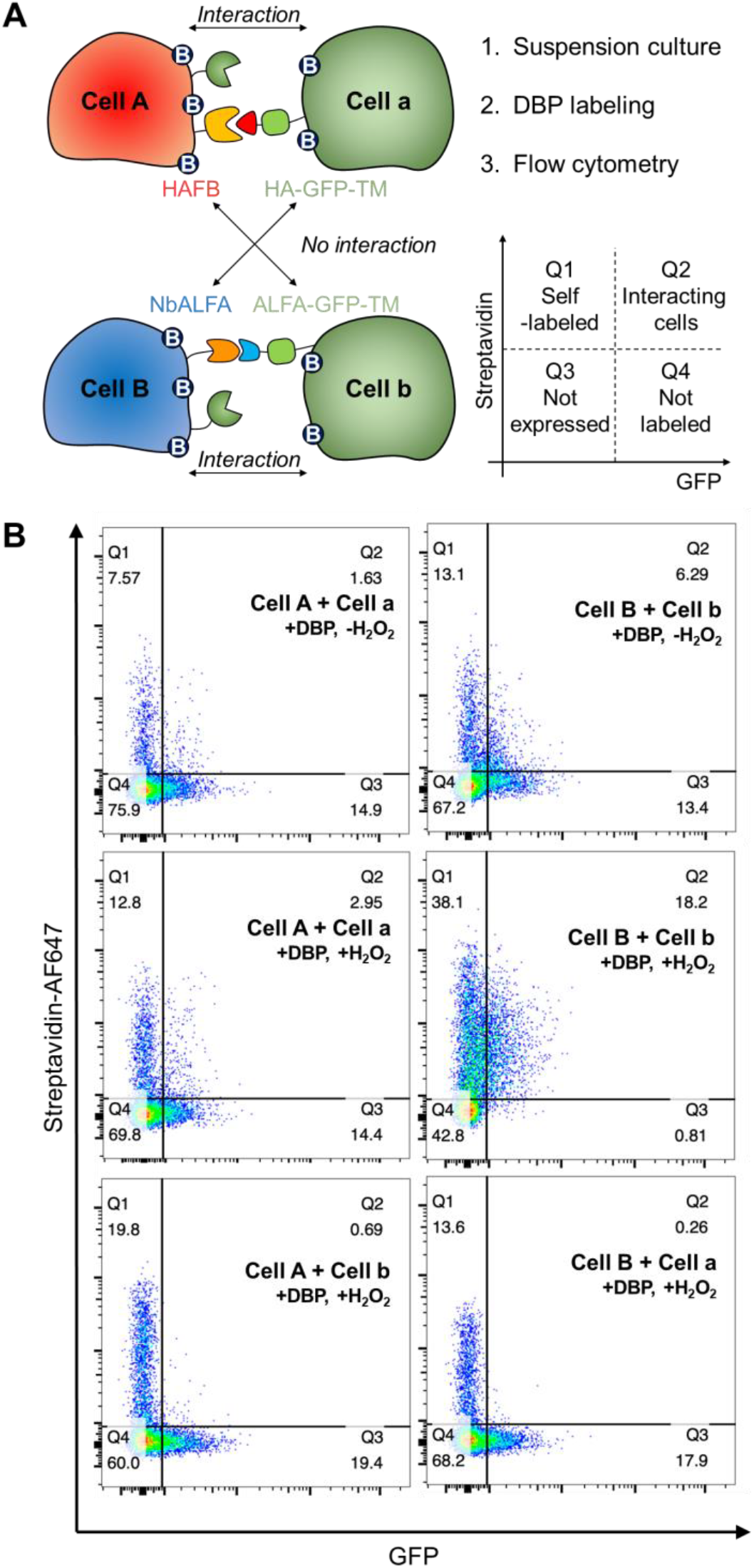
Cell-cell interaction-dependent biotinylation via HRP-TM. (A) Experimental scheme of interaction-dependent biotinylation mediated by HRP-TM. Cell A and Cell a were mixed either with Cell B or Cell b, followed by the sequential addition of DBP. Cells were then stained with streptavidin-AF647 and analyzed by flow cytometry. (B) Flow cytometric analysis of HAFB (αHA)-HA tag and NbALFA (αALFA)-ALFA tag interaction-dependent biotinylation via HRP-TM, even in the absence of exogenous H_2_O_2_.

To assess cell-cell interaction-dependent biotinylation, four groups of cells were prepared. Cell A and Cell B were both transfected with HRP-TM, along with HAFB-TM or NbALFA-TM, respectively. Cell a and Cell b were transfected with 3xHA-GFP-TM or ALFA-GFP-TM, respectively. Cell A and Cell B were then either mixed with Cell a or Cell b, followed by DBP for biotinylation, with or without exogenous H_2_O_2_. The cells were subsequently stained and analyzed by flow cytometry. In this experiment, we also tested the combination of Cell A and Cell b, or Cell B and Cell a, which were expected to have no interaction. In flow cytometry graph, Q1 (Streptavidin positive and GFP negative) represents self-labeled cells by HRP-TM, Q2 (Streptavidin positive and GFP positive) represents interacting proximal cells, Q3 (Streptavidin negative and GFP negative) represents cells that were not expressed, and Q4 (Streptavidin negative and GFP positive) represents HAFB or NbALFA expressed cells but not biotin labeled (**Figure 4A**). In addition, the labeling efficiency was calculated by dividing the fraction of both GFP and streptavidin positive cells by that of total GFP positive cells (**Figure S3**).

Under H_2_O_2_ treatment condition (**Middle, Figure 4B**), both Cell A and Cell B successfully labeled their interacting proximal cells (Q2). Because NbALFA has a lower K_D_ than HAFB, the labeling of proximal cells in the NbALFA-ALFA pair (Cell B - Cell b, 95.7%) was more efficient than in the HAFB-HA pair (Cell A - Cell a, 17.0%). In mismatched pairs (Cell A - Cell b, Cell B - Cell a), almost no biotinylation (3.4%, 1.4%) was observed even under H_2_O_2_ treatment (**Bottom, Figure 4B**), indicating the interaction specificity between two ligand-receptor pairs. Interestingly, matched cells (Cell A - Cell a, Cell B - Cell b) were biotinylated (9.8%, 31.9%) even without exogenous H_2_O_2_ (**Top, Figure 4B**). Consistent with other data, HRP-TM labeled the proximal cells in an interaction-specific manner by utilizing endogenously generated H_2_O_2_.

## Discussion

In this study, we showed that HRP, not APEX2, can facilitate endogenously generated H_2_O_2_ on the cell surface and label proximal cells. Our findings imply that HRP-mediated biotinylation can be used for *in vivo* applications on the cell surface to detect cell-cell networks without toxic H_2_O_2_ treatment. Several studies have used HRP for cell-surface proteome profiling. Li et al. ^[36]^ developed an HRP-based technique for cell-surface proteome profiling in the brain tissue of *Drosophila*. Shuster et al. ^[31]^ applied HRP-mediated cell-surface proteomics to the mouse brain. Since HRP showed no toxicity issue in their expression in live animal model in these studies ^[28, 33]^, we expect that our cell-cell interaction labeling technique can be employed in these HRP-expressed animal models. We believe that HRP-mediated proximal cell labeling has advantages in overcoming the limitations of other approaches, such as utilization of high molecular weight of probe (LIPSTIC, EXCELL, PUP-IT, FucoID) ^[9, 10, 11, 12]^, light irradiation (μMap, PhoTag, PhoXCELL, Ru-^1^O_2_-hydrazide) ^[15, 16, 17, 18]^, low concentration of extracellular ATP (TurboID) ^[37]^, and exogenous treatment of H_2_O_2_ (APEX2).

While utilizing endogenously generated H_2_O_2_ for initiating HRP-mediated labeling offers advantages, there are some caveats associated with this approach. Firstly, the production of H_2_O_2_ can vary across different cell types. Secondly, even within the same cell type, the levels of H_2_O_2_ production may fluctuate under different cellular states. These factors can influence the effectiveness of capturing cell-cell interactions. For example, variations in the quantity of H_2_O_2_ produced by cells before and after drug treatment may lead to changes in the labeling of interacting cells. Such fluctuations may not accurately represent the cell interactions themselves but rather arise from shifts in H_2_O_2_ production levels.

Recently, an integrated approach utilizing proximity labeling and omics data has been reported. Oslund et al. ^[16]^ combined PhoTag with multiomics single-cell sequencing and discovered specific T cell subtype in human peripheral blood mononuclear cell (PBMC) that interacted more with Raji B cells. Zhang et al. ^[38]^ applied QMID to the mouse spleen and characterized gene expression profiles of CD4+ or CD8+ proximal cells by single-cell RNA sequencing (scRNA-Seq). However, these methods are limited to *ex vivo* models, thus prompting increased interest in developing a tool suitable for *in vivo* models. In this context, Hamachi et al. proposed a new bacterial tyrosinase-based PL method, emphasizing its efficient application *in vivo* ^[39]^. In this paper, we propose the possibility that HRP-TM could paint proximal cells in *in vivo* models by utilizing endogenously generated H_2_O_2_. Exploring the interaction between viruses and cells is also an interesting theme, as viruses rely on receptor proteins to invade host cells ^[40]^. By integrating the outcomes of HRP-mediated cell labeling with additional spatial omics data, such as single-cell RNA sequencing (scRNA-seq), we anticipate a comprehensive elucidation of the intricate cell-cell network implicated in disease progression. This approach promises to provide valuable insights into the dynamic cellular landscape underlying various pathological conditions.

## ASSOCIATED CONTENT

### Supporting Information

The Supporting Information is available free of charge at http://pubs.acs.org.

Supporting Information: Materials and methods, construct information, and additional experimental figures including immunofluorescence microscopy images and Western blot (PDF).

## AUTHOR INFORMATION

## Author Contributions

I.J. performed experiments. K.K. and H.W.R. supervised the research. I.J., K.K., and H.W.R. wrote the manuscript.

## Notes

The authors declare no competing financial interest.

## ACKNOWLEDGMENT

This work was supported by the National Research Foundation of Korea (NRF-2022R1A2B5B03001658, NRF-2022M3H9A2096199, NRF-2022M3E5E8081185, RS-2023-00265581 and RS-2023-00260454 to H.W.R, NRF-2022R1C1C2004982 to K.e.K.) and Korea Health Industry Development Institute (KHIDI) funded by the Ministry of Health & Welfare and Ministry of Science and ICT, Republic of Korea (grant numbers: HU23C0204). H.-W.R. was supported by the Samsung Science and Technology Foundation (SSTF-BA2201-08).

## ABBREVIATIONS

HRP: Horseradish peroxidase
TM: Transmembrane domain
DBP: Desthiobiotin-phenol
CMFDA: 5-Chloromethylfluorescein diacetate
GFP: Green fluorescent protein

## Table of Contents Artwork

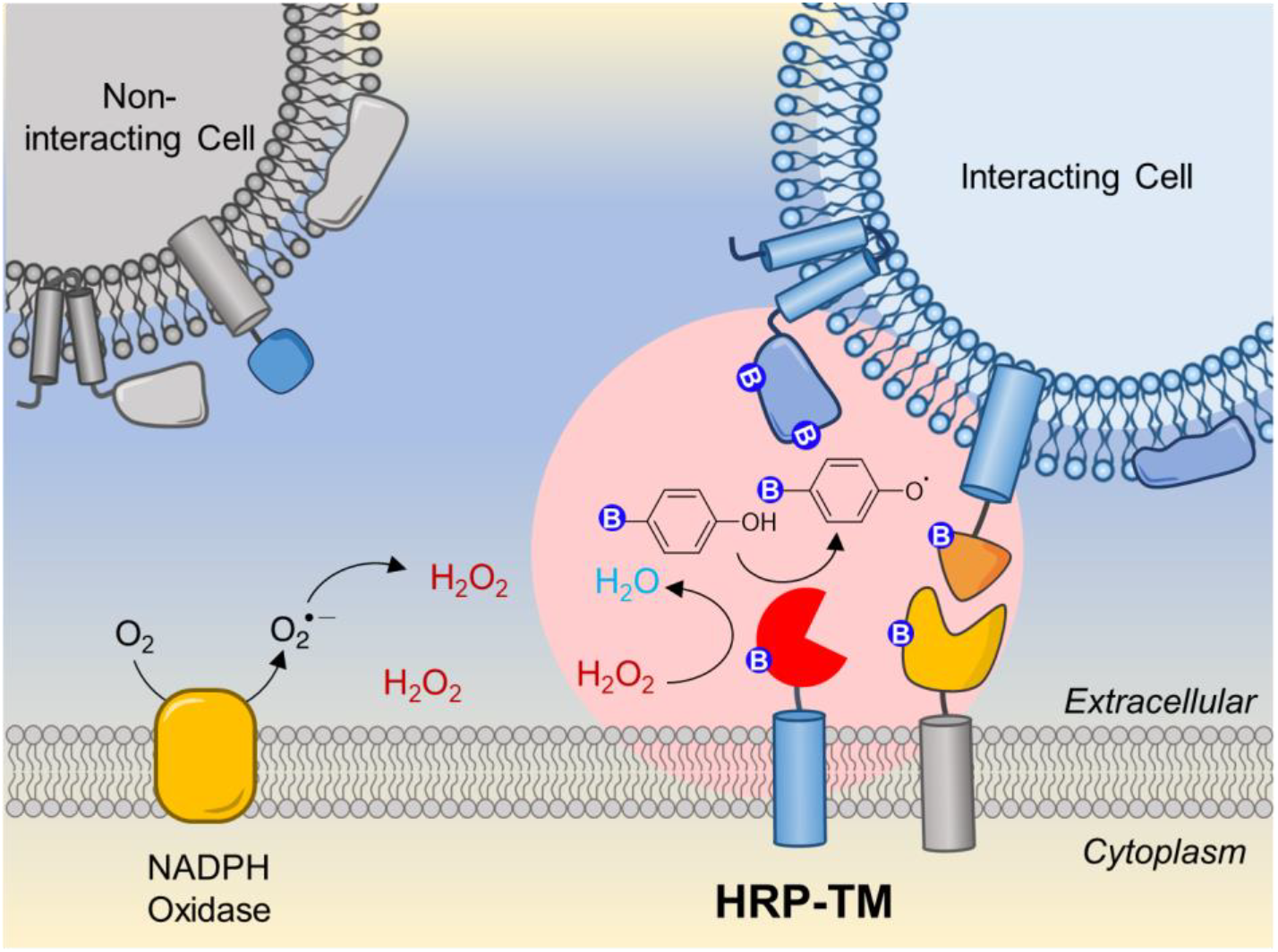

